# Using DNA flow-stretching assay as a tool to validate the tagging of DNA-binding proteins for single-molecule experiments

**DOI:** 10.1101/2023.03.19.533373

**Authors:** Miranda Molina, Lindsey E. Way, Zhongqing Ren, Qin Liao, Xindan Wang, HyeongJun Kim

## Abstract

Due to the enhanced labeling capability of maleimide-based fluorescent probes, lysine-cysteine-lysine (KCK) tags are frequently added to proteins for visualization. In this study, we employed *in vitro* single-molecule DNA flow-stretching assay as a sensitive way to assess the impact of the KCK-tag on the property of DNA-binding proteins. Using *Bacillus subtilis* ParB as an example, we show that, although no noticeable changes were detected by *in vivo* fluorescence imaging and chromatin immunoprecipitation (ChIP) assays, the KCK-tag substantially altered ParB’s DNA compaction rates, its response to nucleotide binding and to the presence of the specific sequence (*parS*) on the DNA. While it is typically assumed that short peptide tags minimally perturb protein function, our results urge researchers to carefully validate the use of tags for protein labeling. Our comprehensive analysis can be expanded and used as a guide to assess the impacts of other tags on DNA-binding proteins in single-molecule assays.

**Motivation:** Single-molecule fluorescence microscopy has been extensively used in modern biology to define the molecular action of proteins. Appending short peptide tags is a common strategy to enhance fluorescence labeling. In this Resources article, we evaluate the impact of a commonly used tag, the lysine-cysteine-lysine (KCK) tag, on protein behavior in single-molecule DNA flow-stretching assay, which is a sensitive and versatile method to understand the action of DNA-binding proteins. Our motivation is to provide researchers with an experimental framework to validate the fluorescently labeled DNA-binding proteins in single-molecule methods.

## Introduction

Fluorescence-based protein visualization has played an instrumental role in single-molecule experiments.^1–5^ Intensive research has led to fruitful development of a variety of fluorophores and tagging methods for proteins.^6–10^ Regardless of the kinds of probes and labeling modalities, the main goals have remained largely unchanged: achieving adequate fluorescence labeling while minimally perturbing the properties of the proteins.

Unwanted alteration of protein properties can occur at any stage during the preparation of the labeled protein, which must be minimized to reveal the true function of the protein of interest. First, introducing additional components into a protein requires a careful consideration during the protein design. The sizes of common fluorescent proteins are around 25 kD,^5, 11^ and those of self-labeling proteins SNAP-, CLIP-, and Halo-tags are about 20 kDa, 20 kDa, and 33 kDa, respectively.^12^ It has been well appreciated that large protein tags could interfere with the protein’s function and disrupt protein-protein interactions, and smaller tags are prefered.^3, 13^ Second, fluorescent-dye-labeling steps can adversely affect protein activities. For example, *Bacillus subtilis* structural maintenance of chromosomes (BsSMC) protein is capable of compacting flow-stretched DNA.^14^ We previously observed that using a centrifugal concentrator-based method to remove unreacted fluorescent dyes abolished BsSMC’s DNA compaction ability whereas using a resin-based column did not.^14^ Third, fluorescent dyes could disrupt the proper functioning of a protein itself. Especially, it has been shown that hydrophobic fluorescent dyes have a potential to cause artifacts due to non-specific binding.^15^ Lastly, fluorescently labeled proteins may malfunction in the context of experimental environments. Prominent examples were reported with quantum dot (QD)-labeled proteins. Effective diameters of commercially available functionalized QDs (“big” QDs) are in the 14-35 nm range^16^ while “small” QDs are 9-12 nm in diameter.^17, 18^ When the localizations of AMPA receptors (AMPAR) on neurons were examined, big QD-labeled AMPARs localized differently from small QD-labeled (or 4 nm organic fluorescent dye-labeled) AMPARs, along with difference in diffusion coefficients, possibly due to the narrow synaptic cleft size (∼30 nm).^19, 20^

While fluorescently tagged proteins are crucial in a variety of biological studies, a challenge is the lack of predictions for how the fluorescent tags or probes will alter protein function. Thus, empirical investigations must follow. The purpose of this study is to explore the single-molecule DNA flow-stretching assay as a sensitive and efficient tool to test whether a tag alters the function of a DNA-binding protein. We chose the DNA flow-stretching approaches because it has been extensively used in studying actions of DNA-binding proteins on individual DNA molecules.^14, 21–26^ Furthermore, contrary to force-based single-molecule assays which visualize one DNA molecule a time, single-molecule DNA flow-stretching assay allows 10-40 DNA molecules to be analyzed in each field of view, which is advantageous for subsequent statistical data analyses.

In this study, for a proof of principle, we tested the effect of lysine-cysteine-lysine (KCK) tag on *B. subtilis* ParB (BsParB) proteins. We chose this tag because maleimide-conjugated fluorescent dyes have been widely used to label proteins via covalent conjugation to surface-exposed cysteines.^1^ However, labeling all desired cysteines with maleimide dyes is not always achieved. The reaction efficiency between the thiol group on cysteine and the maleimide moiety of a fluorescent dye can be increased by flanking the cysteine with two positively charged lysine residues. It was revealed that the neighboring lysine residues decrease pKa of the cysteine residue, thereby increasing thiol-maleimide reactivity.^27–30^ Thus, appending the lysine-cysteine-lysine (KCK) tag to a protein has been a popular and extensively used method due to its superior fluorescence labeling efficiency.^23, 31–37^

We chose ParB protein as our example because of its well known *in vivo* and *in vitro* activities. The ParABS DNA partitioning system is a broadly conserved segregation machinery for bacterial chromosomes and plasmids. ParB binds to *parS* sequences and spreads to neighboring regions^38, 39^ to form a nucleoprotein complex, which is translocated by ParA.^38, 39^ *In vivo*, ParB spreading is evident by two approaches: fluorescence microscopy in which fluorescently-tagged ParB proteins form foci in live cells and chromatin immunoprecipitation (ChIP) assays in which ParB protein associates with 10-20 kb DNA regions encompassing *parS*.^38, 39^ Importantly, it was recently discovered that ParB protein is a novel enzyme that utilizes cytidine triphosphate (CTP) to modulate ParB spreading.^40–42^ *In vitro*, ParB’s spreading has been shown by imaging fluorescently labeled ParB proteins on doubly tethered (or doubly trapped) DNAs with protein loadblocks.^41, 43, 44^ Here we report that DNA compaction by ParB is artificially enhanced by KCK-tags in single-molecule DNA flow-stretching assays *in vitro*. Further investigation indicates that electrostatic interactions between the negatively charged DNA backbone and the positively charged KCK-tag contribute at least partly to these artifacts. Contrary to the *in vitro* single-molecule results, the KCK-tag did not lead to any noticeable changes *in vivo*. In sum, our single-molecule DNA flow-stretching assay is highly sensitive and allows the detection of the property changes in DNA-binding proteins for single-molecule experiments. Its high throughput data production allows statistical analyses and leads to conclusions more efficiently. We propose that the DNA-flow-stretching-based approaches can be used as a tool to detect property changes of DNA-binding proteins upon addition of tags or fluorescent probes.

## Results

### KCK-tags increase BsParB’s DNA compaction rates *in vitro*

KCK tags are frequently used for *in vivo* and *in vitro* protein labels due to its small size and increased labeling efficiency of maleimide-fluorescence dyes.^27–30^ To understand whether this three-amino-acid tag has any impact on ParB proteins, we purified tagged and untagged wild-type *B. subtilis* ParB (BsParB(WT)) proteins (Figure S1) and employed single-molecule DNA flow-stretching assays with a lambda DNA substrate (Figure 1A). Since ParB has been shown to be a CTPase,^40–42^ our samples were treated with apyrase to remove residual nucleotides from our protein samples. Upon addition of the purified proteins, we measured the speed of DNA compaction by tracking the positions of a fluorescent quantum dot labeled at one DNA end (Figure 1B).^23^ In the presence of 50 nM untagged BsParB(WT), we observed robust DNA compaction all the way to the DNA-tether point in the absence of CTP as previously shown (Figures 1B and S2).^23^ Interestingly, both CTP and CTPγS (a non-hydrolyzable CTP analog) inhibited DNA compaction by 39-fold and 149-fold, respectively (Figure 1C), implying counter-productive roles of CTP binding in DNA compaction. The mechanism of CTP binding on ParB’s action is currently being investigated in a separate study. When BsParB(WT) with the KCK-tag at its N-terminus (hereafter “KCK-BsParB(WT)”) (Figure S1) was subjected to the same experiment, without CTP, we observed that the lambda DNA was robustly compacted to the tether point albeit at a slower rate than BsParB(WT) (Figures 1C and S2). However, inclusion of CTP or CTPγS led to strikingly increased DNA compaction rate (10.5-fold and 19.4-fold for CTP and CTPγS, respectively) in KCK-BsParB(WT) compared with untagged BsParB(WT) (Figure 1C). Since in our experience batch-to-batch variations in purified proteins only lead to up to two-fold differences for DNA compaction rates, these dramatic changes prompted us to investigate further.

**Figure 1.**
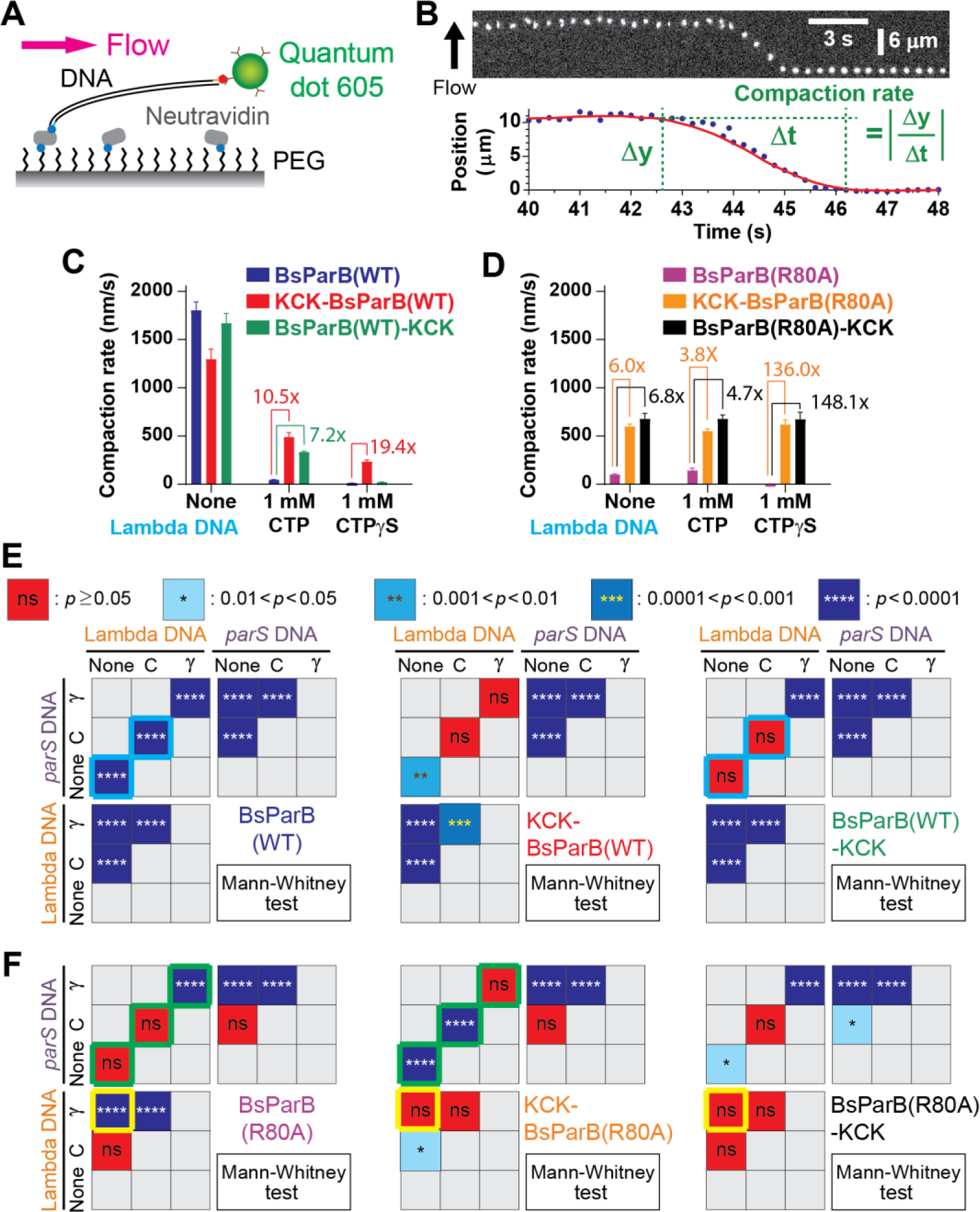
*in vitro* quantitative and qualitative BsParB compaction rate changes by the KCK-tags. (A) Schematic of single-molecule DNA flow-stretching assays. (B) An example of DNA compaction by 50 nM BsParB(WT) protein (top) and the definition of compaction rate (bottom). (C-D) Lambda DNA compaction rates by 50 nM (C) wild-type and (D) R80A mutant proteins. Numbers indicate compaction rate fold increases. Error bars: SEM. (E) Top: The Mann-Whitney test (the Wilcoxon rank sum test) *p*-value color scheme. Bottom: Mann-Whitney test comparisons for compaction rates by wild-type BsParB and its KCK-versions. (F) Mann-Whitney test comparisons for BsParB(R80A) and its KCK-versions. (E-F) Cyan, green, and yellow boxes highlight qualitative protein property changes due to the KCK-tags for visual aids. (C-F) See Tab 1 in the Supplemental File for detailed sample number (*N*) information.

**Figure 2.**
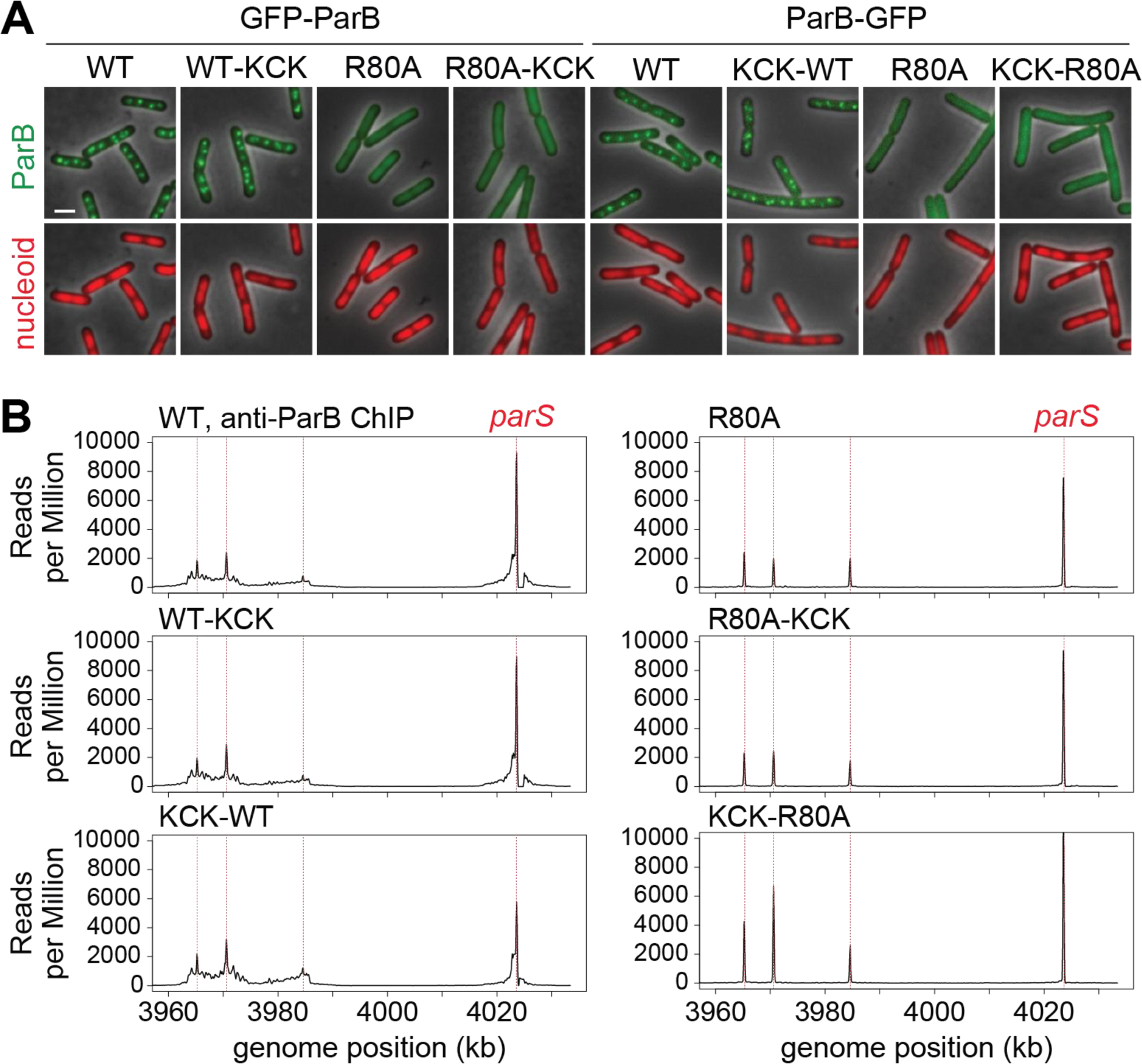
KCK tags do not affect *in vivo* BsParB localization or spreading. (A) Localization of fluorescently tagged ParB(WT) and ParB(R80A) (green). The nucleoid is labeled with HBsu-mCherry (red), and phase-contrast images are shown in gray. Scale bar represents 2 μm. (B) ChIP-seq of wild-type and mutant ParB association with a region of the *B. subtilis* chromosome from 354° to 360° (3960–4033 kb of strain the PY79 genome). Red dotted lines indicate the positions of the four *parS* sites. The number of reads were normalized by the total number of reads per sample. Whereas wild-type ParB spreads several kilobases from *parS* sites, the R80A mutant is restricted to the immediate vicinity of each *parS* site. KCK tags at the N-terminus or C-terminus of wild-type ParB or R80A mutant do not change the property of the variants.

Next, we examined the effect of ParB specific *parS* sequence on DNA compaction rates by inserting a *parS* in the middle of the lambda DNA (hereafter, “*parS* DNA”).^23^. We found that without any nucleotides, untagged BsParB(WT) compacted *parS* DNA 25% slower than it did for lambda DNA without *parS* (Figures S3A and S3C). In the presence of CTP or CTPγS, untagged BsParB(WT)’s compaction rate of *parS* DNA decreased by 16-fold and 50-fold, respectively (Figure S3A). The mechanism of *parS* on ParB’s action is currently being investigated in a separate study. Strikingly, KCK-BsParB(WT) exhibited substantial increase in the *parS* DNA compaction rates in the presence of CTP (4.9-fold) or CTPγS (7.2-fold) compared with untagged BsParB(WT) (Figure S3A). Thus, the KCK-tag enhanced BsParB(WT)’s DNA compaction rate (compared to the untagged BsParB(WT)) when nucleotides are present on both lambda DNA and *parS* DNA.

Given that ParB protein’s CTP binding pocket resides at the N-terminal domain (NTD) and the NTD is implicated to be the DNA-entry gate,^41, 42, 45^ we questioned if the unexpected compaction rate increases also occur when KCK is tagged at the C-terminal of BsParB(WT) protein (hereafter, “BsParB(WT)-KCK”) (Figure S1). Indeed, like KCK-BsParB(WT), BsParB(WT)-KCK also showed much faster compaction with CTP compared with BsParB(WT) (Figure 1C for the lambda DNA and Figure S3A for the *parS* DNA). Thus, KCK enhanced BsParB(WT)’s DNA compaction rate when appended to either terminus.

### KCK-tags alter BsParB’s DNA compaction properties in response to nucleotides and *parS*

To quantify how the BsParB(WT) protein and its KCK-tagged variants respond to different nucleotides and a *parS* site, Mann-Whitney tests were performed for DNA compaction rates with all possible permutations (Figure 1E. Also see Figure S4A). Our analyses revealed that, without any nucleotide or with 1 mM CTP, BsParB(WT) was responsive to the existence of *parS* (*p*<0.001) while BsParB(WT)-KCK did not make statistically significant compaction rate changes with *parS* (*p*≥0.05) (See cyan boxes in Figure 1E). We note that the KCK-tags not only changed compaction rates (Figures 1C, 1D, and S3A-S3C) but also reversed the trend of compaction. Specifically, without nucleotides, when a *parS* site was added to DNA, BsParB(WT)’s compaction rate was slowed down, but KCK-BsParB(WT)’s compaction rate was increased (Figure S3C). These results show that the KCK-tag alters the DNA-compaction ability both quantitatively and qualitatively.

### The KCK-tag alters the action of BsParB R80A mutant

We next investigated whether the compaction rate change induced by the KCK-tag was limited only to the wild-type BsParB. The R80A mutant of BsParB has been shown to abolish proper *in vivo* sporulation, localization, and spreading along with *in vitro* lambda DNA compaction in the absence of nucleotides.^23, 46, 47^ Surprisingly, without nucleotides, although its DNA compaction rate was 18.2-fold lower than BsParB(WT) (Figure S3C), BsParB(R80A) (Figure S1) was still capable of compacting the lambda DNA (Figure 1D), contradicting a previous report (See Discussion.)^23^ Next, we wondered whether a KCK tag alters BsParB(R80A)’s action on DNA. Indeed, with lambda DNA, the compaction rates of both KCK-BsParB(R80A) and BsParB(R80A)-KCK were substantially increased for all tested nucleotides (Figure 1D). When *parS* DNA was used as a substrate, compaction rate increases by KCK tags (*p*<0.0001) were also noted (Figure S3B). The visualized Mann-Whitney comparison charts for DNA compaction rates highlight that BsParB(R80A), KCK-BsParB(R80A), and BsParB(R80A)-KCK respond differently to different nucleotides and the presence of *parS* (See green and yellow boxes in Figure 1F. Also see Figure S4B.)

### The effects of KCK-tags on protein action are limited to *in vitro* assays but not *in vivo*

The different effects of KCK tags in DNA compaction *in vitro* prompted us to systematically test the effect of KCK tag on BsParB’s or BsParB(R80A)’s localization and spreading *in vivo*. We first generated eight GFP fusions to the ParB variants with KCK tags at the C- or N-terminus of the protein and performed fluorescence microscopy (Figure 2A). Consistent with previous findings that R80A abolishes ParB spreading,^23^ BsParB(WT) formed foci in the cells, while BsParB(R80A) had diffused localization on the DNA. Interestingly, KCK tags at the C- or N-terminus did not alter the localization of ParB(WT) or ParB(R80A) (Figure 2A). In a complementary approach, we analyzed the *in vivo* spreading of ParB variants on the genome by chromatin immunoprecipitation (ChIP-seq) assays using anti-ParB antibodies (Figure 2B). We observed that BsParB(WT) spreads to a ∼20 kb region surrounding the *parS* site, but BsParB(R80A) did not spread. These results are consistent with previously published data.^23^ Importantly, having a KCK tag at the C- or N-terminus did not affect the spreading of BsParB(WT) or BsParB(R80A). We also show that the KCK-tagged proteins have similar expression levels compared to the matched untagged controls (Figures S5A and S5B). These experiments demonstrate that the KCK tag does not affect BsParB’s functions *in vivo*. Thus, the effects of KCK tags on BsParB(WT) and BsParB(R80A) are specific to *in vitro* experiments.

### Charges on the KCK-tag contribute to the *in vitro* protein property changes

This finding prompted us to understand the mechanism by which the KCK-tag boosts the DNA compaction rate of ParB protein *in vitro*. One possibility for the compaction rate increase is that more BsParB proteins were recruited onto DNA due to interactions between the positively charged KCK-tag and the negatively charged DNA backbone. Alternatively, the KCK-tag could impact the subsequent action of the BsParB proteins while the level of the initial protein recruitment is intact. To distinguish these two possibilities, we directly visualized the recruitment of untagged and KCK-tagged BsParB(R80A) proteins onto lambda DNA. Proteins were nonspecifically labeled with the NHS-ester version of Cyanine3 fluorescent dye, and the moment of the very first labeled protein’s arrival into the camera’s field-of-view was evident by increase in background intensity (Figures 3A and 3B). In this approach, background-subtracted integrated fluorescence intensity on DNA is directly proportional to the amount of BsParB protein recruited onto the DNA. The microscopy showed that the background-subtracted integrated fluorescence intensities with KCK-BsParB(R80A) and BsParB(R80A)-KCK were higher than those with BsParB(R80A) (*p*<0.0001) (Figure 3C). Thus, our data show that the KCK-tags enhanced protein loading and increased compaction rates with a caveat that our experimental approaches do not address if the KCK tags impact on subsequent protein action after being recruited onto DNA.

**Figure 3.**
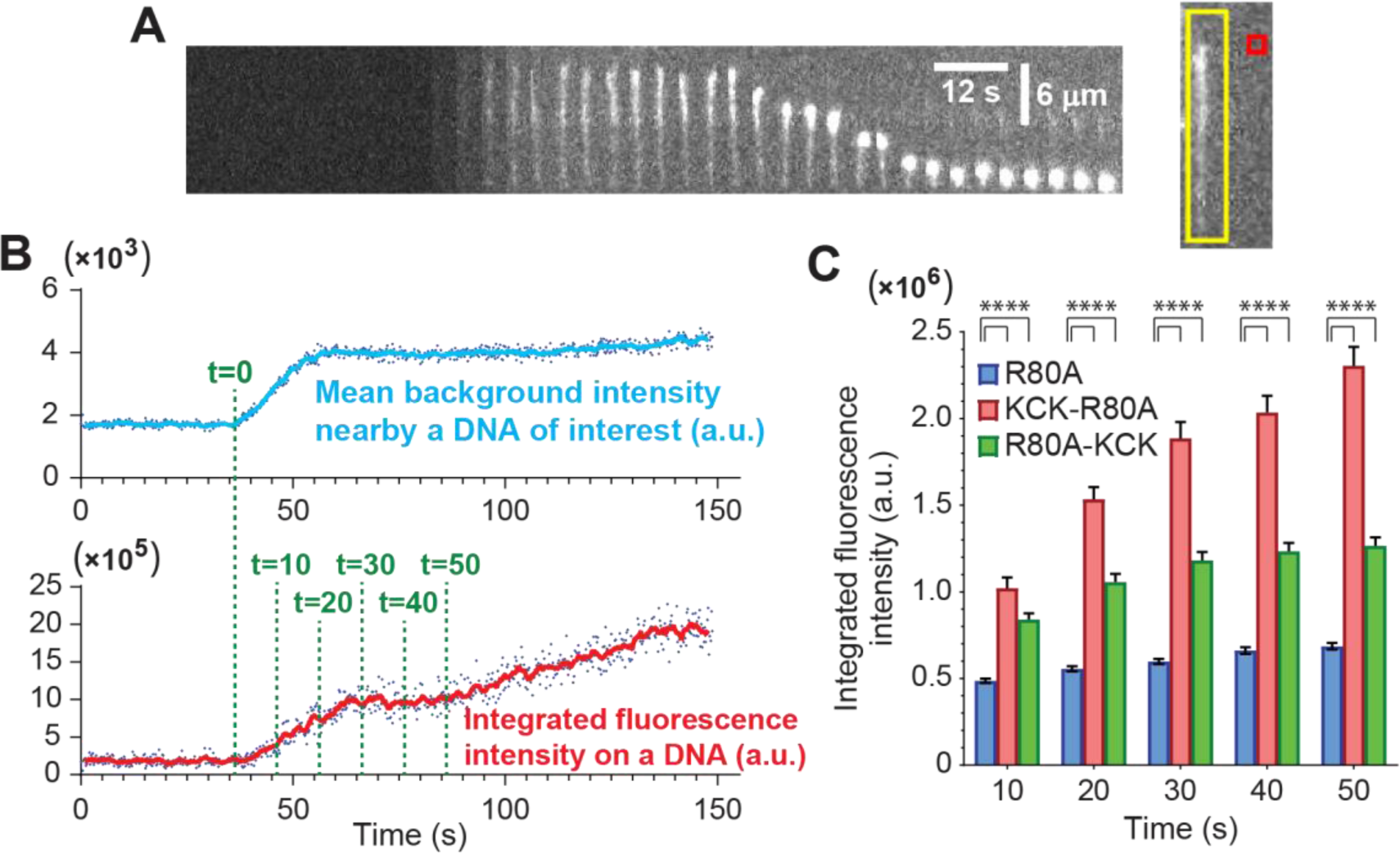
More BsParB(R80A) proteins are loaded onto flow-stretched lambda DNAs when the KCK-tag is appended. (A) (Left) A representative kymograph for DNA flow-stretching experiments with fluorescently-labeled BsParB protein. 30 nM KCK-BsParB(R80A) without any nucleotides. (Right) The mean background intensity was obtained for the area bound by the red square (0.96×0.96 μm), and the background-subtracted integrated fluorescence intensity was obtained for the area bound by the yellow box (2.4×14.4 μm). (B) Time-trajectories for mean background and integrated fluorescence intensities on the DNA shown in (A). The time point when the mean background intensity starts to increase is defined as t=0. DNA flow-stretching experiments were performed with fluorescently-labeled proteins. (C) Integrated fluorescence intensities on lambda DNAs by cyanine3-labeled BsParB(R80A) (*N*=51), KCK-BsParB(R80A) (*N*=40), and BsParB(R80A)-KCK (*N*=26) measured at different time points. Error bars: s.e.m., **** denotes *p*<0.0001. See Tab 1 in the Supplemental File for detailed sample number (*N*) information.

To address whether the charge of KCK tags was the issue, we prepared recombinant wild-type and R80A mutant BsParB proteins where a negatively charged glutamic acid-cysteine-glutamic acid (ECE) tag was added to the N-terminus of proteins. If electrostatic interactions between the appended tags and DNA backbone contribute to *in vitro* artifacts, slower compaction rates are expected with ECE-tagged BsParB proteins (hereafter “ECE-BsParB”) due to repulsive forces between negative charges. As expected, DNA compactions by ECE-BsParB(R80A) were noticeably inefficient. The compaction rates by ECE-BsParB(R80A) are significantly lower (0.001<*p*<0.01 and *p*<0.0001) than those by BsParB(R80A) regardless of the presence of the *parS* DNA sequence and CTP (Figure 4A). Consistent with this observation, the ECE-BsParB(WT) protein also exhibits inefficient DNA compaction compared with its BsParB(WT) counterpart in the absence of any nucleotides (Figure S6).

**Figure 4.**
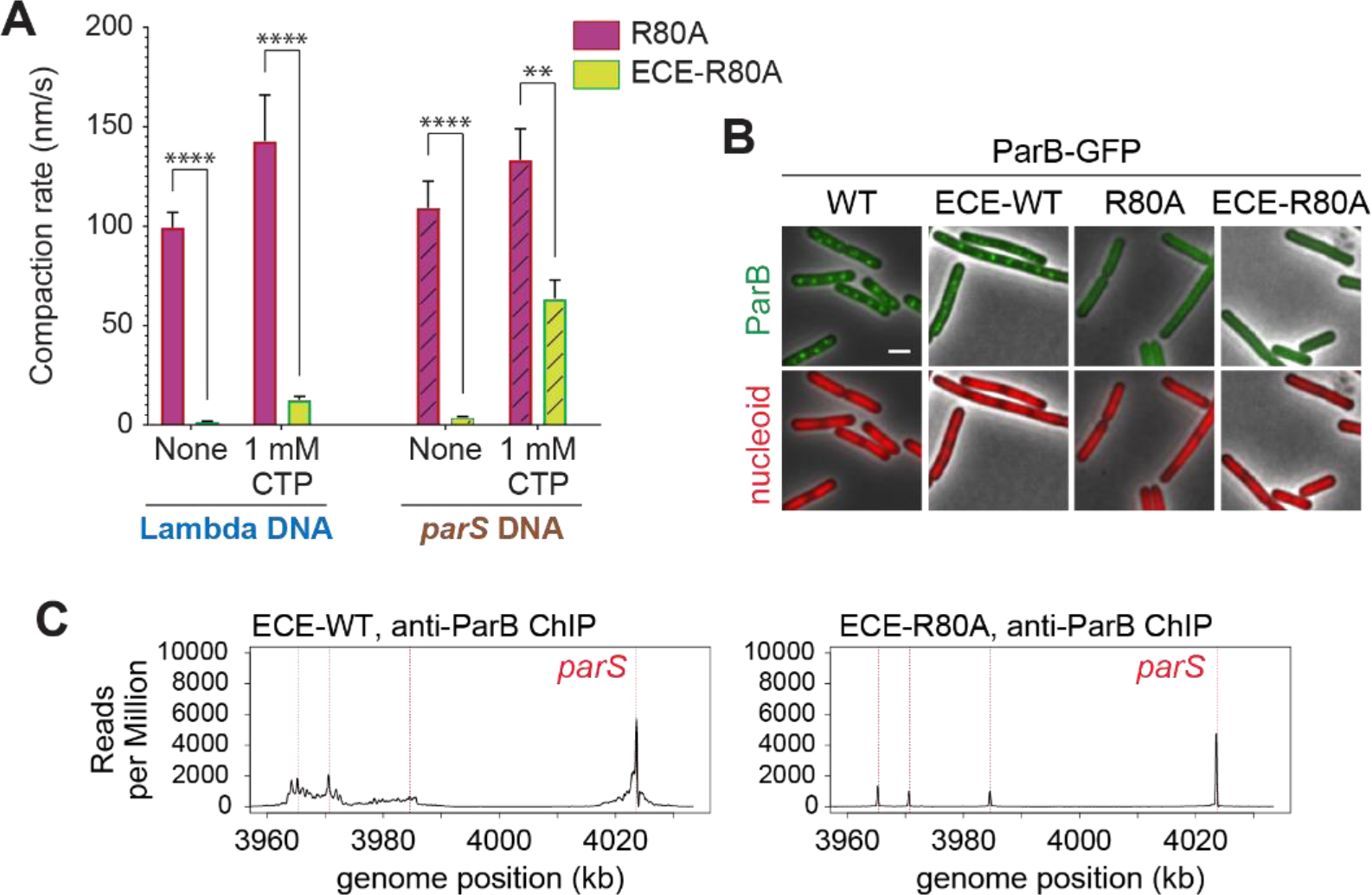
Adding negatively charged ECE-tag on BsParB protein slows down DNA compaction in *vitro* but does not lead to *in vivo* changes. (A) Lambda and *parS* DNA compaction rates by BsParB(R80A) and ECE-BsParB(R80A) both in the presence and absence of CTP. Error bars: s.e.m., ** and **** denote 0.001<*p*<0.01 and *p*<0.0001, respectively. See Tab 1 in the Supplemental File for detailed sample number (*N*) information. (B) Localization of fluorescently tagged BsParB(WT), BsParB(R80A), and their ECE-tagged versions (green). Red: the nucleoid labeled with Hbsu-mCherry. Gray: phase-contrast images. Scale bar represents 2 μm. (C) ChIP-seq of ECE-tagged wild-type (left) and mutant ParB (right) association with a region of the *B. subtilis* chromosome from 354° to 360° (3960–4033 kb of strain the PY79 genome). Red dotted lines indicate the positions of the four *parS* sites. The number of reads were normalized by the total number of reads per sample. Whereas ECE-ParB(WT) spreads several kilobases from *parS* sites, the ECE-R80A mutant is restricted to the immediate vicinity of each *parS* site.

Next, we investigated any *in vivo* property changes caused by N-terminally appended ECE-tag. Fluorescence microscopy experiments show that the ECE-tag does neither abolish the *in vivo* fluorescence foci formation with the wild-type BsParB protein nor lead to the formation of clear foci with the R80A mutant BsParB (Figure 4B). Additionally, ChIP-seq assays using anti-ParB antibodies indicate that wild-type BsParB proteins spread to a ∼20 kb regions around the *parS* site and the R80A mutant does not spread regardless of the presence of the ECE-tag (Figure 4C). All *in vivo* results consistently demonstrate that the KCK and ECE tags appended to BsParB proteins do not have noticeable impacts. The effects of the tags are only limited to *in vitro* assays, and electrostatic interactions between charged residues on the tag and the DNA backbone are at least partly responsible for the *in vitro* effects.

## Discussion

In this study, we employed single-molecule DNA flow-stretching assay to measure DNA compaction rates. Contrary to traditional ensemble measurements, single-molecule approaches can reveal unprecedentedly detailed information by looking into individual molecules and providing statistical analyses. Using BsParB protein as an example, we were able to detect the effect of small tags on protein action at high sensitivity across a wide range of conditions.

Fluorescent labeling of BsParB proteins had been mainly achieved by either replacing an amino acid residue with cysteine (such as S68C^43, 48^ and L5C^44^) or appending the KCK-tag.^23^ The previous study showed that KCK-tagged BsParB protein did not lead to disruption on *in vivo* GFP-BsParB foci formation and BsParB-dependent SMC complex (SMC-ScpA-ScpB) localizations.^23^ Furthermore, DNA motion capture assay showed that Cy3-labeling of BsParB through the KCK-tag did not alter end-biased lambda DNA compaction patterns using *in vitro* assays.^23^ However, our thorough assessments of untagged and the KCK-tagged BsParB proteins by the single-molecule DNA flow-stretching assay revealed unexpected alternations in the protein recruitment level and DNA compaction rates.

The increased compaction rates shown with the KCK-tagged BsParB proteins are ascribed, at least partly, to electrostatic interactions between opposite charges on the KCK-tag and DNA backbone as evidenced by the enhanced fluorescence intensity on DNA and results of the ECE-tagged BsParB. However, we note that the KCK-tag affected not only quantitative compaction rates but also qualitative behaviors of the protein against different nucleotide statuses and the presence of a *parS* sequence. Furthermore, in the absence of CTP, the lambda DNA compaction stopped before reaching the tether point with BsParB(WT)-KCK protein while BsParB(WT) and KCK-BsParB(WT) proteins compacted DNA all the way to the tether point (Figure S2). Therefore, it is possible that KCK-tags affect BsParB protein actions after its loading onto DNAs.

Previously, the R80A mutant of BsParB protein has been shown to be incapable of compacting DNA.^23, 43, 46^ However, slow but robust DNA compaction by BsParB(R80A) protein was observed in this study. Although both studies used the same assay, in our study, we supplemented magnesium ions to our buffer as a cofactor of CTP and used apyrase during our protein purification to remove residual CTPs. Since BsParB(R80A) is deficient in CTP hydrolysis^41^, it is possible that CTPs were co-purified with BsParB(R80A) in the previous study^23^. Consistent with our speculation, in the absence of Mg^2+^ and the presence of CTP, BsParB(R80A)’s compaction rate was reduced dramatically (Figure S7), providing an explanation for the undetectable compaction by BsParB(R80A) in the previous study.

Although only the KCK-tagged BsParB proteins were tested with single-molecule DNA flow-stretching assay, our approach could be readily applicable to other DNA-binding proteins. For proteins capable of compacting DNA molecules, the DNA compaction rates of proteins containing desired tags (such as Halo-, CLIP-, SNAP-, KCK-, and sortase-tags) need to be measured. DNA motion-capture assay (a variant of the single-molecule DNA flow-stretching)^14, 21–23, 49^ can be also performed, and the comparison between untagged- and tagged-proteins will help determine the validity of the tagged protein. Once intact or minimally perturbed activity of the tagged proteins is confirmed, those tags can be labeled with a desired fluorescent dye, and the compaction rates can be measured again to detect any changes. In this case, the fluorescent dye on the protein and the quantum dot at the end of the DNA must be spectrally distinct to allow the compaction rate measurements.^14, 22^ For DNA-binding proteins incapable of compacting DNAs, useful information can still be obtained by measuring and comparing integrated fluorescence intensities from untagged- and tagged-proteins.

In summary, deep understanding of any biological system requires both *in vitro* and *in vivo* approaches. Our study reveals that addition of short peptide tags may produce misleading *in vitro* results despite normal functionality *in vivo.* Additionally, our results raise a possibility that fluorescent dyes conjugated to a DNA-binding protein result in altered *in vitro* protein activities due to electrostatic interactions between charges on the fluorescent probe and those on the DNA backbone. Whenever adding a small amino acid tag is desired for *in vitro* experiments, careful controls must be performed to ensure that this does not perturb the activity of the protein. Here we show that this highly sensitive single-molecule DNA flow-stretching assay can be used as a powerful and efficient validation tool for modified proteins.

### Limitation of the study

Here using KCK-tag on BsParB protein as a model system, we evaluate single-molecule DNA flow-stretching assay as an efficient and sensitive tool to assess the impact of small tags on protein action. While this study focuses on the methodology to validate protein tagging, the mechanism of BsParB action is being investigated in a separate study. Interestingly, although the KCK-tags drastically altered ParB’s action at *in vitro* single-molecule level, they did not change ParB’s activity in *B. subtilis* live cells in any detectable way. These results imply that other cellular factors, such as the property of the cytoplasm and/or the biophysical aspects of the chromatin, make BsParB more tolerant to tagging than that in the minimalist *in vitro* setups.

Here we have discovered and evaluated that small protein tags can lead to the change of protein property in single molecule experiments. However, the scope of this study is limited to only BsParB protein and the KCK-tags. It is desirable that other DNA-binding proteins with different fluorophore labeling platforms be investigated for future studies, such as site-specific labeling methodologies involving unnatural amino acids (UAAs)^13, 50, 51^ or aldehyde tags^52^. Furthermore, assessing whether a charged fluorescent probe that labels the KCK-tag can rescue the changes caused by the tag alone will be an interesting topic for future studies.

Finally, this DNA flow-stretching assay is applicable to proteins that have reasonable affinity to DNA. If the protein’s DNA-binding affinity is too low, the fluorescence background might mask the signal from the DNA-bound proteins.

## Acknowledgements

We thank the Kim and Wang labs for discussions and support, Xheni Karaboja for assisting with ChIP-seq and microscopy experiments, Patrick Sheets for assisting with strain building, Candice Etson for providing the *E. coli* strain containing *parS*-lambda DNA, Thomas Graham for providing plasmids for BsParB wild-type and R80A mutant, Alan Grossman for strains and antibodies, and the Indiana University Center for Genomics and Bioinformatics for high throughput sequencing. We thank Thomas Graham for reading the manuscript and providing insightful feedback. Support for this work comes from National Institutes of Health R01GM141242, R01GM143182 (X.W.), and R35GM143093 (H.K.).

## Author contributions

M.M. and H.K. purified proteins, performed single-molecule experiments, and analyzed data. L.E.W. constructed plasmids and strains, performed microscopy, and immunoblot analysis. Z.R., Q.L., and X.W. performed ChIP-seq and analysis. X.W. designed, analyzed, and supervised the *in vivo* experiments. H.K. designed, analyzed data, and supervised the *in vitro* experiments. M.M. and L.E.W. contributed to writing the method sections of the paper. X.W. and H.K. wrote the paper with input from all authors.

## Methods

### KEY RESOURCES TABLE

- Please refer to the Supplemental File.

### RESOURCE AVAILABILITY

#### Lead contact

- Further information and requests for resources should be directed to and will be fulfilled by the lead contact, HyeongJun Kim.

#### Materials availability

- This study did not generate new unique reagents. Plasmids generated in this study are available from the corresponding authors upon request.

#### Data and code availability

- A list of figures that have associated raw data can be found from Tabs 6 and 9 in the Supplemental File. Single-molecule analysis data can be found in Tabs 1-5 in the Supplemental File.
- The datasets generated during and/or analyzed during the current study are available from the corresponding authors on reasonable request.
- The Matlab codes used in single-molecule data are available from our previous publication^14^. Alternatively, the codes will be available from the corresponding author (H.K.) upon request.

## EXPERIMENTAL MODEL AND SUBJECT DETAILS

### Plasmid constructions for *in vitro* single-molecule assays

Plasmids harboring coding sequences of His6-SUMO-BsParB(WT) (pTG011)^23^, His6-SUMO-KCK-BsParB(WT) (pTG042)^23^, His6-SUMO-BsParB(R80A) (pTG037)^23^, and His6-SUMO-KCK-BsParB(R80A) (pTG044)^23^ were generous gifts from Thomas Graham. Site-directed mutagenesis were performed to generate plasmids harboring coding sequences of His6-SUMO-BsParB(WT)-KCK (m0067) and His6-SUMO-BsParB(R80A)-KCK (m0069) using oHK050F and oHK050R as primers. The plasmid harboring coding sequences of His6-SUMO-ECE-BsParB(WT) (m0064) were generated using oHK048F and oHK048R as primers and m0043 as a substrate. Contrary to other plasmids, the plasmid harboring coding sequences of His6-SUMO-ECE-BsParB(R80A) (m0070) was generated by following the vendor-supplied NEBuilder HiFi DNA Assembly Master Mix (NEB E2621S, Ipswich, MA) protocol. First, the His6-SUMO-BsParB(WT) plasmid (pTG011 = m0041) was linearized and the majority of SUMO-BsParB(WT) coding sequences were removed by PCR using oHK038F and oHK038R as primers. Then, gfHK009 and gfHK010 were used as gene fragments with both containing 23 bp overlaps. After NEBuilder HiFi DNA assembly, NEB 5-alpha competent *E. coli* cells (NEB C2987H, Ipswich, MA) were transformed with the reaction mixture. The sequences were confirmed using T7, oHK023, oHK024, oHK025, and oHK026 oligos. See Tabs 7-8 in the Supplemental File for their sequences.

### Protein expression and purification

Rosetta2(DE3)pLysS competent cells (EMD Millipore, Burlington, MA) transformed with a plasmid were cultured overnight at 37°C in the presence of 100 μg/mL ampicillin and 20 μg/mL chloramphenicol. 1 L of LB medium with 80 μg/mL ampicillin was inoculated with the overnight culture and grown at 37°C until the OD_600_ reached 0.4-0.6. Protein expression was induced with 500 μM isopropyl-β-D-thiogalactoside (IPTG), and the culture was shaken at 30°C for an additional 4 hours. The cells were harvested by centrifugation at 4°C. The cell pellets were resuspended in PBS buffer and spun at 5,000 g. They were resuspended in ParB lysis buffer (20 mM Tris, pH 8.0, 1 M NaCl, 50 mM imidazole, 5 mM 2-mercaptoethanol), supplemented with 0.1 mM phenylmethylsulfonyl fluoride (PMSF) and a protease inhibitor cocktail (Roche, Basel, Switzerland) (Total volume: 45 mL), and flash-frozen. BsParB proteins were purified based on a two-step tandem purification method as previously described^23^ but with some modifications. Briefly, after thawing the harvested cells, additional 0.9 mM PMSF (total 1.0 mM PMSF), 50 mg/mL lysozyme, 3 μL of universal nuclease (Thermo Fisher Scientific 88701, Waltham, MA), and 5 mM 2-mercaptoethanol were added, and it was left in ice for 30 minutes. Cells were lysed by sonification and centrifuged twice in an FA-6×50 rotor: first at 11,000 g for 30 minutes, then at 20,133 g for 30 minutes. The clarified supernatant was incubated with Ni-NTA agarose beads (Qiagen, Hilden, Germany) for 1 hour in the presence of 1 unit of apyrase (NEB, Ipswich, MA) and 5 mM MgCl_2_, to help minimize cellular NTPs that may otherwise be co-purified, and 1 tablet of cOmplete Mini EDTA-free protease inhibitor cocktail (Roche, Basel, Switzerland). The Ni-NTA agarose resin was washed with lysis buffer (supplemented with 5 mM MgCl_2_) followed by ParB salt-reduction buffer (20 mM Tris, pH 8.0, 350 mM NaCl, 50 mM imidazole, 5 mM MgCl_2_, 5 mM 2-mercaptoethanol). The proteins were manually eluted ten times with 1.5 mL of ParB elution buffer (20 mM Tris, pH 8.0, 350 mM NaCl, 250 mM imidazole, 5 mM MgCl_2_, 5 mM 2-mercaptoethanol).

The peak fractions of ParB protein were pooled and treated with His6-Ulp1 protease to remove the N-terminal His6-SUMO tag^23^. The pooled proteins and His6-Ulp1 protease were dialyzed together overnight at 4°C against ParB dialysis/storage 1 buffer (20 mM Tris, pH 8.0, 350 mM NaCl, 10 mM imidazole, 5 mM 2-mercaptoethanol, 1 mM MgCl_2_, 10% glycerol). After centrifuging the dialyzed proteins at maximum speed for 10 minutes, the supernatant was allowed to interact with the Ni-NTA resin for at least 1 hour at 4°C. Then, the flowthrough was collected. 0.5 mL of the ParB dialysis/storage 1 buffer to the Ni-NTA resin column was added multiple times, and the eluents were collected. Running an SDS-polyarcylamide (SDS-PAGE) gel indicated that the flowthrough and the eluent fractions contained ParB protein, while the cleaved His6-SUMO and His6-Ulp1 remained in the resin. The flowthrough and the peak fractions were pooled and dialyzed against ParB dialysis/storage 2 buffer (20 mM Tris, pH 8.0, 350 mM NaCl, 10% glycerol), where 5 mM 2-mercaptoethanol was included in case of KCK-tagged protein purifications. The protein concentration was measured by a NanoDrop One spectrophotometer (Thermo Scientific, Waltham, MA) using 32.58 (kDa) and 7,450 (M^−1^ cm^−1^) as its molecular weight and extinction coefficient, respectively. The purified proteins (Figure S1) were run on a precast polyacrylamide gel (Bio-Rad, Hercules, CA) with Tris/Glycine/SDS running buffer (Bio-Rad, Hercules, CA). InstantBlue Coomassie protein stain (Abcam, Cambridge, United Kingdom) was used to stain for the polyacrylamide gel. The gel image was obtained using UVP UVsolo touch gel documentation system (Analytik Jena, Jena, Germany) and provided in the Figure S1 without any image processing.

### DNA and Quantum-dot preparations

One end of bacteriophage lambda DNA (or *parS* DNA^23^) was labeled with a biotin to tether the DNA onto the single-molecule microfluidic flowcell, and the other end was labeled with a digoxigenin to attach a quantum dot (Figure 1A) as previously described^14, 22^. Briefly, Lambda-BL1Biotin and Lambda-Dig2 oligos (Tab 8 in the Supplemental File) were treated with T4 polynucleotide kinase (PNK) (NEB, Ipswich, MA) for phosphorylation at 37°C for 1 hour. A 15-fold molar excess of the phosphorylated Lambda-BL1Biotin oligo was introduced for annealing to a 12-base 5’ single-stranded overhang on one end of lambda DNA (or *parS* DNA^23^). The mixture of DNA and oligo was incubated at 65°C for 10 minutes and slowly cooled down, and then ligated by T4 ligase for 2 hours at room temperature. The other end of the lambda DNA (or *parS* DNA) was tagged with a digoxigenin by supplementing a 60-fold molar excess of the phosphorylated Lambda-Dig2 oligo at 45°C. After 30-minute incubation, the mixture was slowly back to room temperature followed by a 2-hour ligation step at room temperature. Since the sequences of Lambda-BL1Biotin and Lambda-Dig2 oligos are complementary to each other, it is important to remove unreacted excess oligos. After running a 0.4% agarose gel overnight at 4°C, the desired DNA band was excised and put into a dialysis tube. Applying an electric field allowed DNAs to leave the excised agarose gel, but DNAs were confined to the dialysis tube volume. DNAs were collected, and ethanol precipitation was performed to recover doubly-tagged lambda DNAs (or *parS* DNAs) in EB buffer (10 mM Tris-Cl, pH 8.5).

As we previously did^14, 21, 22^, anti-digoxigenin antibody-conjugated quantum dot 605 (Invitrogen, Waltham, MA) was prepared following Invitrogen’s Qdot 605 antibody conjugation kit (Q22001MP) manual. However, since this kit was discontinued, all the kit components were separately purchased including Qdot 605 ITK amino (PEG) quantum dots (Invitrogen Q21501MP). For the antibody, anti-digoxigenin fab fragments (Roche 11214667001, Basel, Switzerland) were used.

### BsParB protein labeling with fluorescent dyes

BsParB(R80A) proteins were incubated with sulfo-Cyanine3 NHS ester dye (Lumiprobe 11320, Hunt Valley, MD) at 4°C overnight. Labeled protein was separated from free dye using Micro Bio-Spin P-6 gel columns (Bio-Rad 7326221, Hercules, CA). Each labeled protein and Cyanine3 dye concentrations were measured three times using Nanodrop, and the averaged values were used as final concentrations. The protein labeling efficiencies were 30.1%, 32.0%, and 30.0% for BsParB(R80A), KCK-BsParB(R80A), and BsParB(R80A)-KCK, respectively. These numbers correspond to about 0.6 Cyanine3 dyes per each BsParB protein dimer.

### Single-molecule flow-stretching assays

Surface-passivated coverglasses were prepared by aminopropyl silanization and PEGylation (PEG: polyethylene glycol) as previously described^14, 22^. A microfluidic flow cell was constructed from a quartz plate (Technical Glass Product, Paineville, OH) adhered to the PEGylated coverglass via double-sided tape (Grace Bio-Labs, Bend, OR) with rectangular cuts that make up the flow cell channels. Inlet and outlet tubing were inserted through holes on the quartz plate and made air-tight with epoxy^14, 22^. In-depth description of single-molecule flow-stretching assays was already provided in previous publications^14, 21, 22^. Briefly, about 4% of the PEG on the surface-passivated coverglass contains biotins that serve as a neutravidin binding platform. Pre-mixed quantum dot-labeled biotinylated lambda DNA (or *parS* DNA) was introduced to allow the DNA surface tethering. For experiments with labeled proteins, quantum dot incubation with biotinylated DNA is omitted. After washing unbound DNAs and quantum dots, an intended concentration of BsParB protein was flowed in (with and without nucleotides). Unless otherwise stated, the buffer composition was 10 mM Tris, pH 7.5, 100 mM NaCl, and 2.5 mM MgCl_2_. For the experiments without magnesium ions, the 2.5 mM MgCl_2_ was omitted. CTPγS was custom-synthesized (Jena Bioscience, Jena, Germany). The single-molecule imaging was performed on a semi-custom microscope with a 532-nm laser (Coherent, Santa Clara, CA) built upon the IX-83 total internal reflection fluorescence (TIRF) microscope (Evident Scientific, Olympus, Waltham, MA). The images were recorded every 200 milliseconds with 100-millisecond exposure time using the Micro-Manager software^53^. Regions-of-interest (ROIs) of DNA compaction events were determined using ImageJ (FIJI) software, and the positions of quantum dots as a function of time were determined by Gaussian-fitting-based custom-written Matlab software codes^14^. The compaction rate measurements were taken from distinct samples (quantum dot-bound DNAs).

### Bacterial strains and growth

*B. subtilis* strains were derived from the prototrophic strain PY79^54^. Cells were grown in defined rich Casein Hydrolysate (CH) medium^55^ at 37°C. Strain, plasmids, oligonucleotides, and next-generation sequencing samples used in this study can be found in Tabs 6-9 in the Supplemental File.

### Fluorescence microscopy

Fluorescence microscopy was performed using a Nikon Ti2 microscope (Nikon Instruments, Melville, NY) equipped with Plan Apo 100x/1.45NA phase contrast oil objective and an sCMOS camera. Images were cropped and adjusted using MetaMorph software. Final figure preparation was performed in Adobe Illustrator.

### ChIP-seq

Chromatin immunoprecipitation (ChIP) was performed as described previously^56, 57^. Briefly, cells were crosslinked using 3% formaldehyde for 30 min at room temperature and then quenched using 125 mM glycine, washed using PBS, and lysed using lysozyme. Crosslinked chromatin was sheared to an average size of 250 bp by sonication using Qsonica Q800R2 water bath sonicator. The lysate was precleared using Protein A magnetic beads (GE Healthcare/Cytiva 28951378, Marlborough, MA) and was then incubated with anti-ParB antibodies^58^ overnight at 4°C. The next day, the lysate was incubated with Protein A magnetic beads for 1h at 4°C. After washes and elution, the immunoprecipitate was incubated at 65°C overnight to reverse the crosslinks. The DNA was further treated with RNaseA, Proteinase K, extracted with PCI, resuspended in 100 µl EB and used for library preparation with the NEBNext Ultra II kit (E7645). The library was sequenced using Illumina NextSeq500 (Illumina, San Diego, CA) at IU Center for Genomics and Bioinformatics. The sequencing reads were mapped to *B. subtilis* PY79 genome (NCBI Reference Sequence NC_022898.1) using CLC Genomics Workbench (Qiagen, Hilden, Germany). We note that the genome coordinate of this genome is shifted compared to the *B. subtilis* 168 genome (NC000964) used in our previous study^23^. Sequencing reads were normalized by the total number of reads, plotted and analyzed using R.

### Immunoblot analysis

Exponentially growing cells were collected and resuspended in lysis buffer (20 mM Tris pH 7.0, 1 mM EDTA, 10 mM MgCl_2_, 1 mg/ml lysozyme, 10 µg/ml DNase I, 100 µg/ml RNase A, 1 mM PMSF and 1% proteinase inhibitor cocktail (Sigma-Aldrich P-8340, St. Louis, MO) to a final OD_600_ of 10 for equivalent loading. The cell resuspensions were incubated at 37°C for 10 min for lysozyme treatment, followed by the addition of an equal volume of 2x Laemmli Sample Buffer (Bio-Rad 1610737, Hercules, CA) containing 10% β-Mercaptoethanol. Samples were heated for 15 min at 65°C prior to loading. Proteins were separated by precast 4-20% polyacrylamide gradient gels (Bio-Rad 4561096, Hercules, CA) and electroblotted onto mini PVDF membranes using Bio-Rad Transblot Turbo system and reagents (Bio-Rad 1704156, Hercules, CA). The membranes were blocked in 5% nonfat milk in phosphate-buffered saline (PBS) with 0.5% Tween-20, then probed with anti-ParB (1:5000)^58^ or anti-SigA (1:10,000)^59^ diluted into 3% BSA in 1x PBS-0.05% Tween-20. Primary antibodies were detected using Immun-Star horseradish peroxidase-conjugated goat anti-rabbit antibodies (Bio-Rad 1705046, Hercules, CA) and Western Lightning Plus ECL chemiluminescence reagents as described by the manufacturer (Perkin Elmer NEL1034001, Waltham, MA). The signal was captured using ProteinSimple Fluorchem R system. The intensity of the bands was quantified using ProteinSimple AlphaView software.

### Plasmid construction for *in vivo* experiments

**pWX1092** [*pelB::Psoj-spo0J(ΔparS)-mgfpmut3 tet*] was constructed by an isothermal assembly reaction containing three fragments: 1) pWX516 digested with HindIII and BamHI, and gel purified; 2) *spo0J (ΔparS)* amplified from pWX563^23^ using oWX2974 and oWX2975; 3) *mgfpmut3* amplified from pWX563^23^ using oWX2976 and oWX2977. pWX516 contains *pelB::Psoj (tet)*. The construct was sequenced using oWX507, oWX669, and oWX670.

**pWX1093** [*pelB::Psoj-KCK-spo0J(ΔparS)-mgfpmut3 tet*] was constructed by an isothermal assembly reaction containing three fragments: 1) pWX516 digested with HindIII and BamHI, and gel purified; 2) *KCK-spo0J (ΔparS)* amplified from pWX563^23^ using oWX2978 and oWX2975; 3) *mgfpmut3* amplified from pWX563^23^ using oWX2976 and oWX2977. pWX516 contains *pelB::Psoj (tet)*. The construct was sequenced using oWX507, oWX669, and oWX670.

**pWX1103** [*pelB::Psoj-mgfpmut3-spo0J-R80A(ΔparS)-KCK cat*] was constructed by an isothermal assembly reaction containing two PCR products: 1) pWX611 amplified using oWX3001 and oWX418; 2) pWX611 amplified using oWX3002 and oWX2071. This procedure introduced the R80A mutation to pWX611^23^, which is *pelB::Psoj-mgfpmut3-spo0J(ΔparS)-KCK cat*. The construct was sequenced using oWX507, oWX669, and oWX670.

**pWX1104** [*pelB::Psoj-spo0J-R80A(ΔparS)-mgfpmut3 tet*] was constructed by an isothermal assembly reaction containing two PCR products: 1) pWX1092 amplified using oWX3001 and oWX418; 2) pWX1092 amplified using oWX3002 and oWX2071. This procedure introduced the R80A mutation to pWX1092. The construct was sequenced using oWX507, oWX669, and oWX670.

**pWX1105** [*pelB::Psoj-KCK-spo0J-R80A(ΔparS)-mgfpmut3 tet*] was constructed by an isothermal assembly reaction containing two PCR products: 1.) pWX1093 amplified using oWX3001 and oWX418; 2) pWX1093 amplified using oWX3002 and oWX2071. This procedure introduced the R80A mutation to pWX1093. The construct was sequenced using oWX507, oWX669, and oWX670.

**pWX1106** [*pelB::Psoj-soj-spo0J-R80A(ΔparS)-KCK cat*] was constructed by an isothermal assembly reaction containing two PCR products: 1) pWX612 amplified using oWX3001 and oWX418; 2) pWX612 amplified using oWX3002 and oWX2071. This procedure introduced the R80A mutation to pWX612^23^, which is *pelB::Psoj-soj-spo0J(ΔparS)-KCK cat*. The construct was sequenced using oWX507, oWX1086, and oML77.

**pWX1107** [*pelB::Psoj-KCK-spo0J(ΔparS) tet*] was constructed by an isothermal assembly reaction containing two PCR products: 1) pWX1093 amplified using oWX3004 and oWX418; 2) pWX1093 amplified using oWX3003 and oWX2071. This procedure introduced a stop codon and removed *mgfpmut3* from pWX1093. The construct was sequenced using oWX507 and oML85.

**pWX1108** [*pelB::Psoj-KCK-spo0J-R80A(ΔparS) tet*] was constructed by an isothermal assembly reaction containing two PCR products: 1) pWX1107 amplified using oWX3001 and oWX418; 2) pWX1107 amplified using oWX3002 and oWX2071. This procedure introduced the R80A mutation to pWX1107. The construct was sequenced using oWX507 and oML85.

**pWX1167** *[pelB::Psoj-ECE-spo0J(ΔparS) tet]* was constructed by an isothermal assembly reaction containing two PCR products: 1) pWX1107 amplified using oWX3197 and oWX418; 2) pWX1107 amplified using oWX3198 and oWX2071. This procedure introduced the ECE tag and removed the KCK tag from pWX1107. The construct was sequenced using oWX507 and oML85.

**pWX1168** *[pelB::Psoj-ECE-spo0J-R80A(ΔparS) tet]* was constructed by an isothermal assembly reaction containing two PCR products: 1) pWX1108 amplified using oWX3197 and oWX418; 2) pWX1108 amplified using oWX3198 and oWX2071. This procedure introduced the ECE tag and removed the KCK tag from pWX1108. The construct was sequenced using oWX507 and oML85.

**pWX1169** *[pelB::Psoj-ECE-spo0J(ΔparS)-mgfpmut3 tet]* was constructed by an isothermal assembly reaction containing two PCR products: 1) pWX1093 amplified using oWX3197 and oWX418; 2) pWX1093 amplified using oWX3198 and oWX2071. This procedure introduced the ECE tag and removed the KCK tag from pWX1093. The construct was sequenced using oWX507, oWX669, and oWX670.

**pWX1170** *[pelB::Psoj-ECE-spo0J-R80A(ΔparS)-mgfpmut3 tet]* was constructed by an isothermal assembly reaction containing two PCR products: 1) pWX1105 amplified using oWX3197 and oWX418; 2) pWX1105 amplified using oWX3198 and oWX2071. This procedure introduced the ECE tag and removed the KCK tag from pWX1105. The construct was sequenced using oWX507, oWX669, and oWX670.

### Strain construction

*B. subtilis* strains were generated by successive transformations of plasmids or genomic DNA.

### Statistics and reproducibility

Not all measurement groups passed the normality test (See Tab 3 in the Supplemental File). Therefore, in this study, we report the results of nonparametric Mann-Whitney test in Figures 1E and 1F. However, we obtained similar results from two-sided Welch’s t-test (Figures S4A and S4B) since the t-test results are still valid when the sample sizes are large (>25) and there are not extreme outliers^60^. All the statistical analyses (Shapiro-Wilk normality test, Mann-Whitney test, and two-sided Welch’s t-test due to different variances and sample sizes) for DNA compaction rates were performed using Prism software (GraphPad, San Diego, CA). The exact sample sizes (*N*), mean, and standard error of the mean are provided in Tabs 1 and 2 in the Supplemental File. The normality test results are available in Tab 3 in the Supplemental File. Tabs 4 and 5 in the Supplemental File show the exact *p*-values for comparing wild-type (and its KCK-tagged versions) and R80A mutant (and its KCK-tagged versions) compaction rates, respectively. The reproducibility of single-molecule experiments for each experimental condition was checked by performing the same experiments at least three times.

---

**Figure S1.**
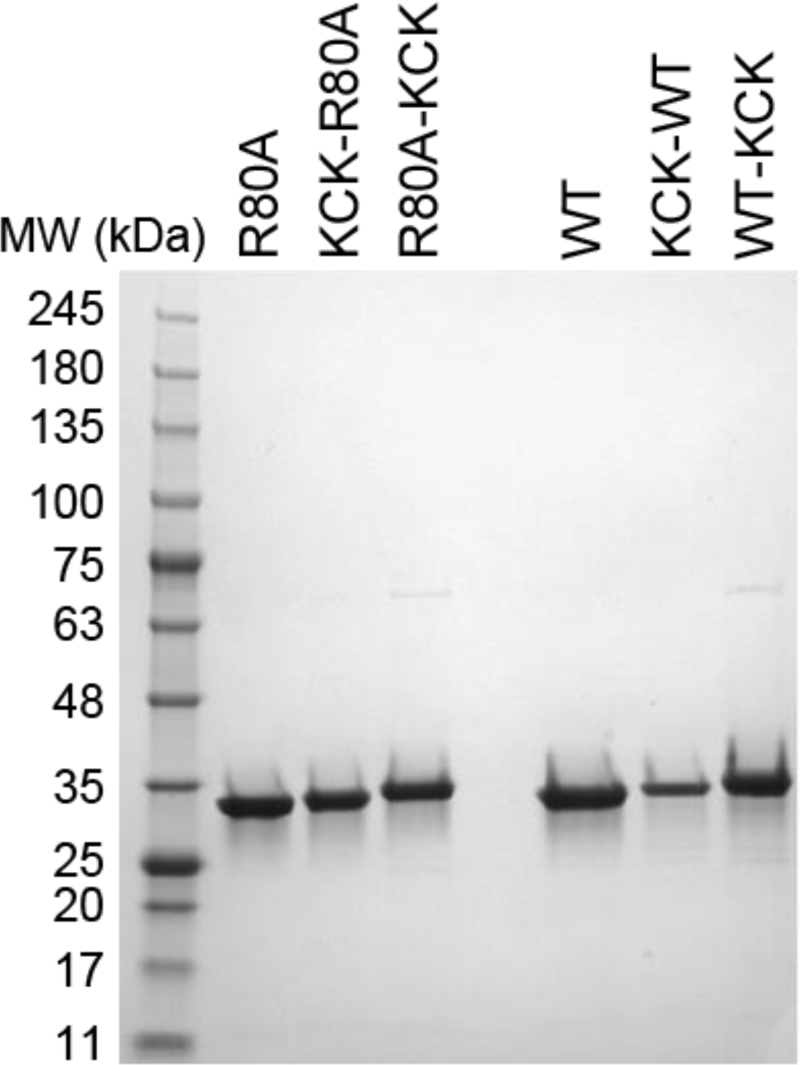
Related to Figures 1, S3, and S7. Purified BsParB proteins. InstantBlue Coomassie-stained SDS-PAGE gel of recombinant *B. subtilis* ParB proteins used in this study. The protein ladder (Gold Biotechnology, #P007-500) is on the left.

**Figure S2.**
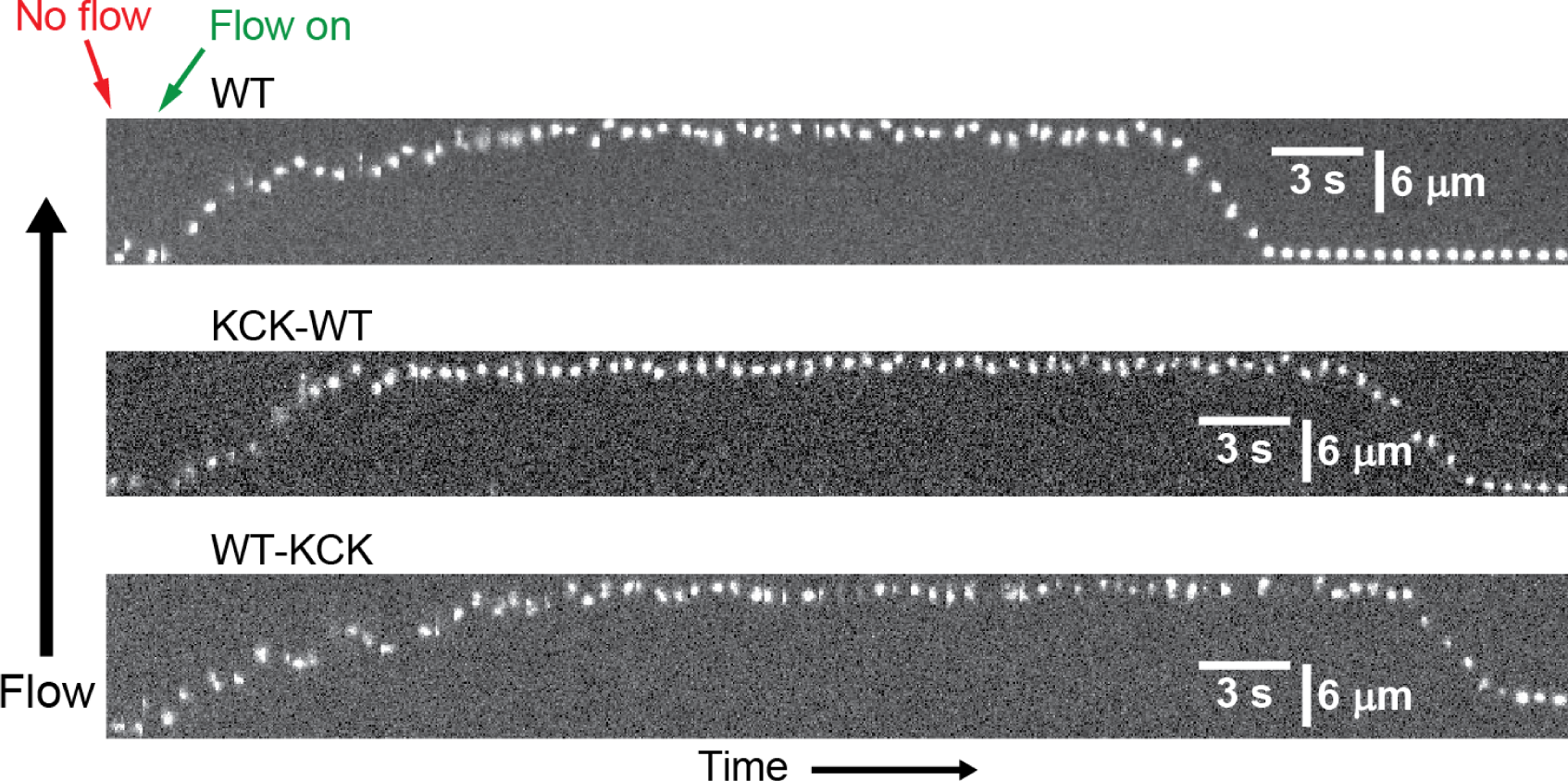
Related to Figure 1. Wild-type BsParB proteins compact DNA towards the tether point. Representative kymographs for (untagged) wild-type BsParB, KCK-BsParB, and BsParB-KCK proteins. One end of DNA is tethered to the microfluidic sample chamber surface, and the other is labeled with a quantum dot 605. In the beginning, the average quantum dot position is approximately the same as the DNA tether point due to the lack of flow. Turning on the flow stretches DNA. Accompanying proteins leads to the DNA compaction towards the tether point. [Protein] = 50 nM, [CTP] = 0, lambda DNA without any *parS* sites.

**Figure S3.**
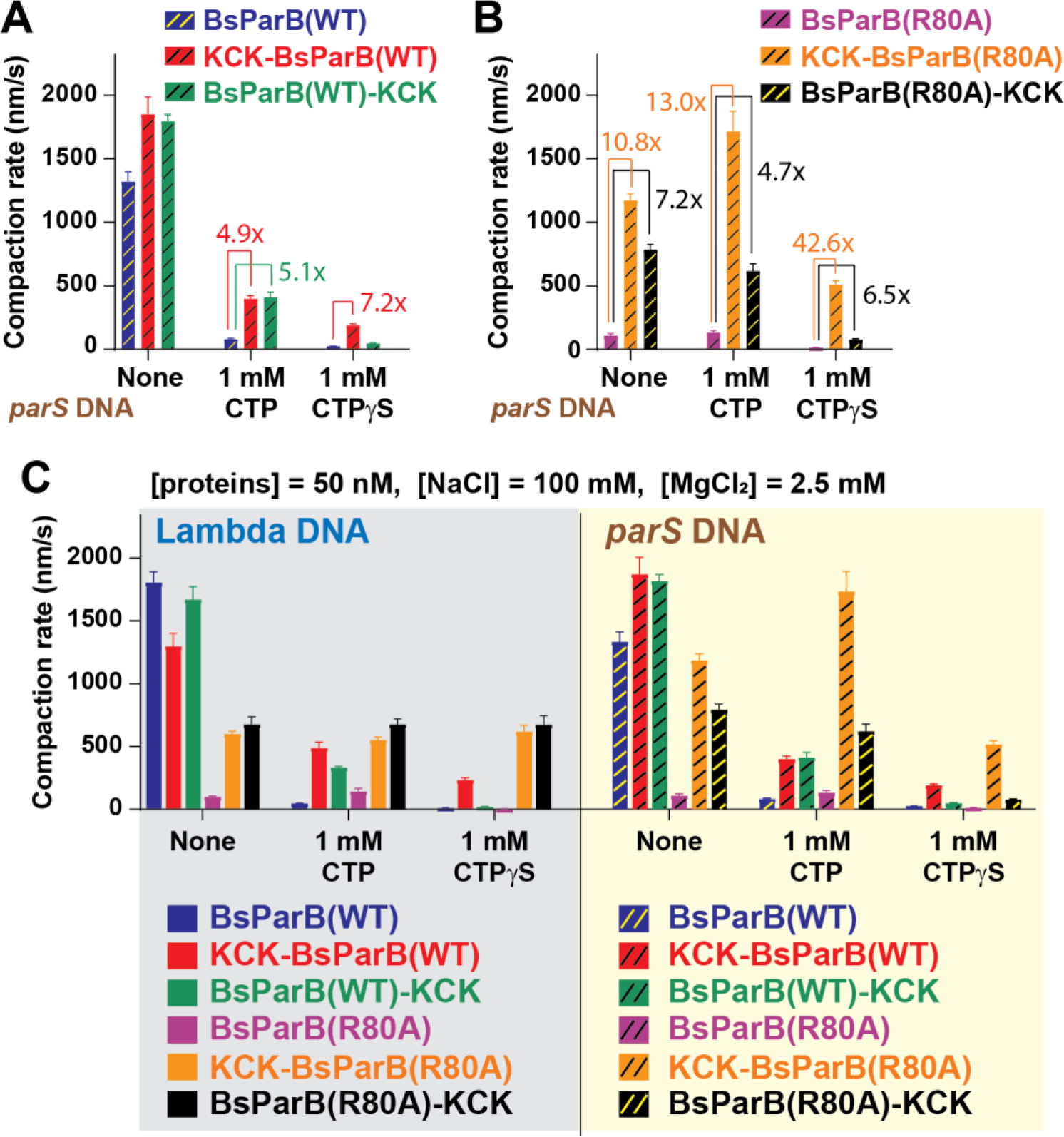
Related to Figure 1. KCK-tags lead to quantitative and qualitative compaction rate changes. (A) *parS* DNA compaction rates by BsParB(WT), KCK-BsParB(WT), and BsParB(WT)-KCK proteins in the absence and presence of 1 mM CTP or 1 mM CTPγS. Error bars: s.e.m., The numbers indicate compaction rate fold changes. (B) *parS* DNA compaction rates by BsParB(R80A), KCK-BsParB(R80A), and BsParB(R80A)-KCK proteins in the absence and presence of 1 mM CTP or 1 mM CTPγS. Error bars: s.e.m., The numbers indicate compaction rate fold changes. (C) For direct comparisons, the compaction rates shown in (A), (B) and Figures 1C and 1D are consolidated. Error bars: s.e.m. (A-C) See Tab 1 in the Supplemental File for detailed sample number (*N*) information.

**Figure S4.**
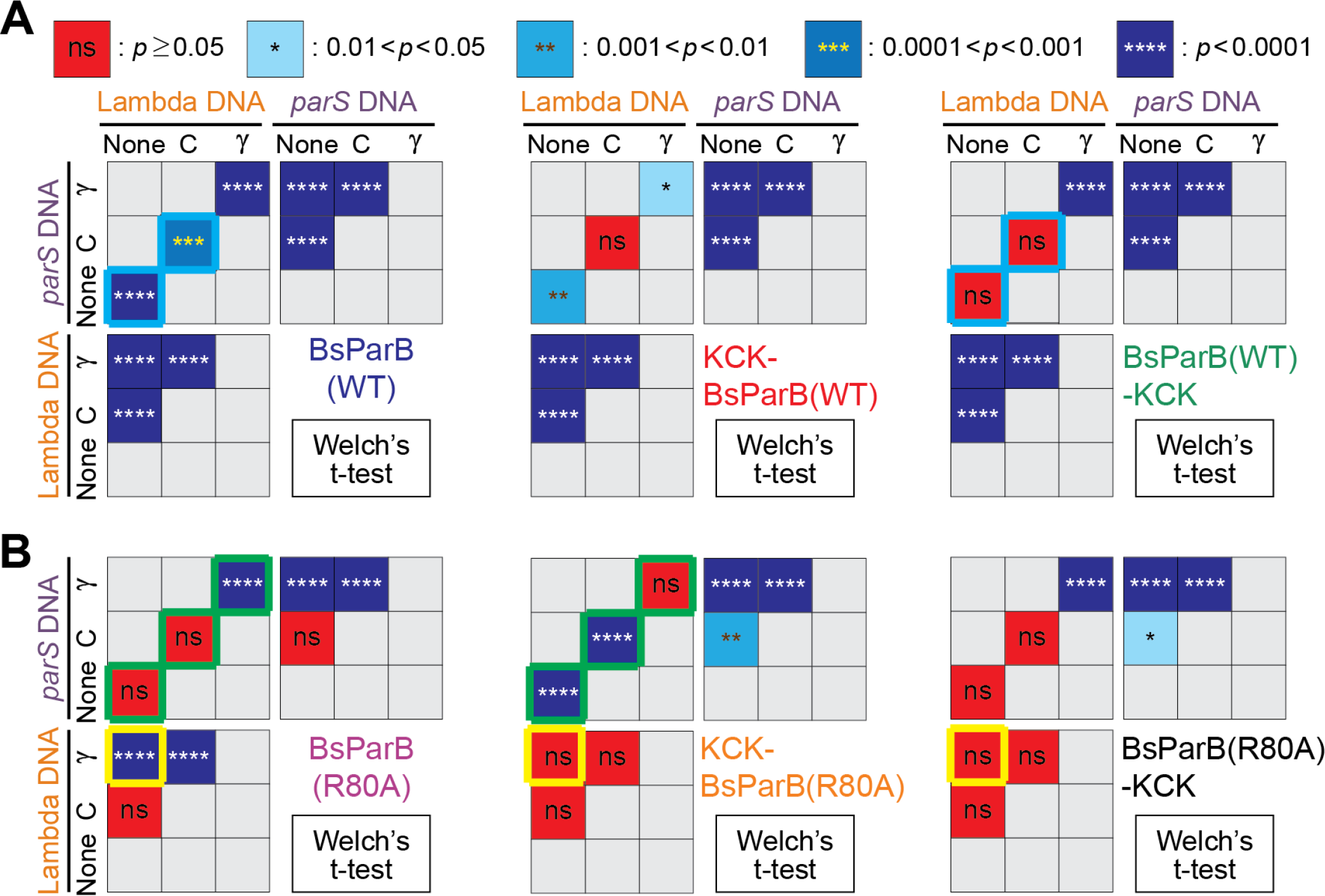
Related to Figure 1. KCK-tags lead to qualitative compaction rate changes. Since not all results pass the Shapiro-Wilk normality test, we employed the Mann-Whitney tests to compare DNA compaction rates in Figures 1E and 1F. However, Welch’s t-test results are still informative as long as there are not extreme outliers and there are enough (>25) data points.60 Indeed, the Welch’s t-test results provided here are very similar to the ones from the Mann-Whitney tests. (A) Top: The Welch’s t-test *p*-value color scheme. Bottom: The Welch’s t-test comparisons for compaction rates by wild-type BsParB and its KCK-versions. (B) The Welch’s t-test comparisons for BsParB(R80A) and its KCK-versions. (A-B) Cyan, green and yellow boxes highlight qualitative protein property changes due to the KCK-tags for visual aids. See Tab 1 in the Supplemental File for detailed sample number (*N*) information.

**Figure S5.**
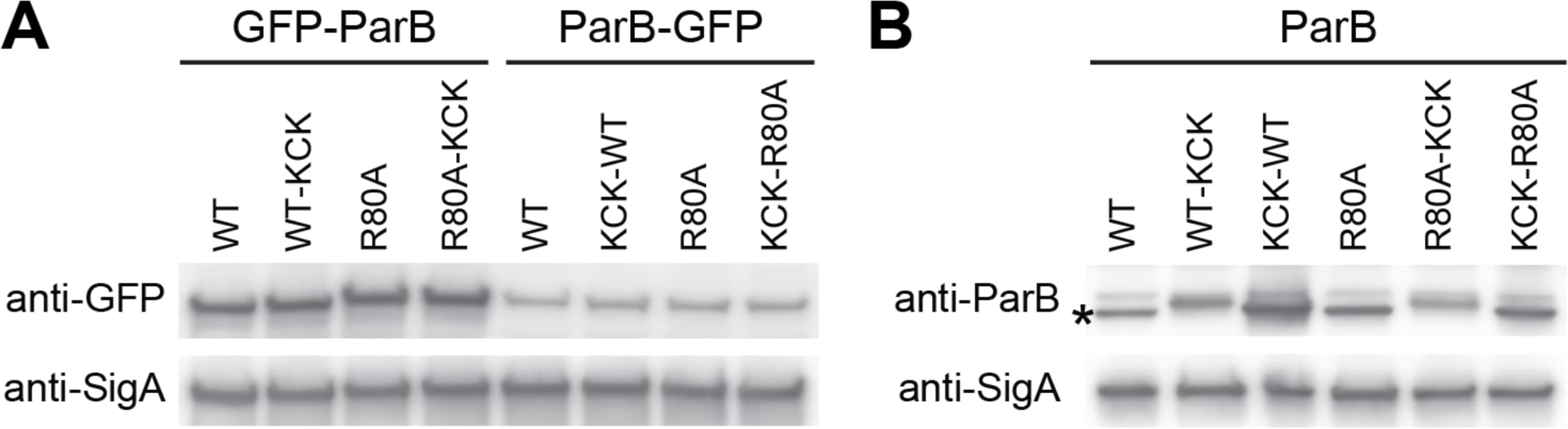
Related to Figure 2. KCK tags do not significantly alter the level of ParB or R80A. (A) Western blot of GFP-tagged ParB variants. Although GFP-ParB levels are higher than ParB-GFP levels, the R80A mutation or KCK tag does not change the protein levels. SigA levels are shown to control for loading. (B) Western blot of ParB variants. The R80A mutation or KCK tag does not dramatically change the protein levels. Asterisk indicates the ParB band. SigA levels are shown to control for loading.

**Figure S6.**
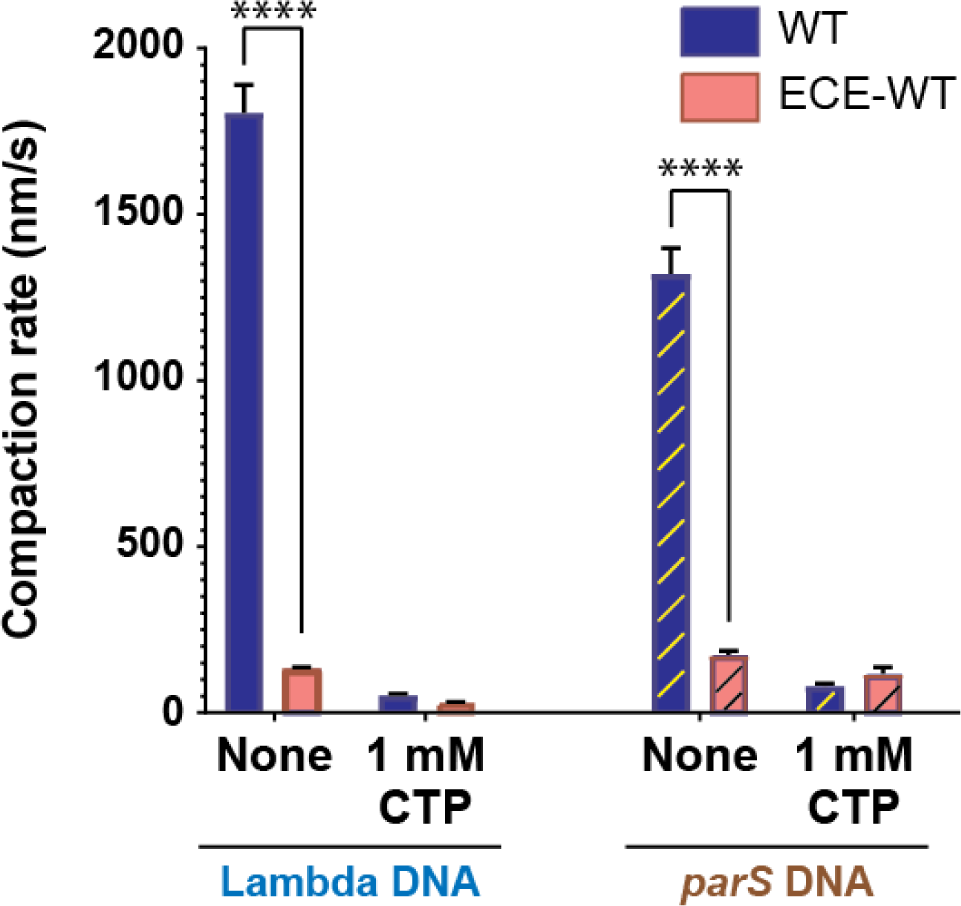
Related to Figure 4. The ECE-tag slows down DNA compaction by the wild-type BsParB protein. Lambda and *parS* DNA compaction rates by BsParB(WT) and ECE-BsParB(WT) both in the presence and absence of CTP. Error bars: s.e.m., **** denotes *p*<0.0001. See Tab 1 in the Supplemental File for detailed sample number (*N*) information.

**Figure S7.**
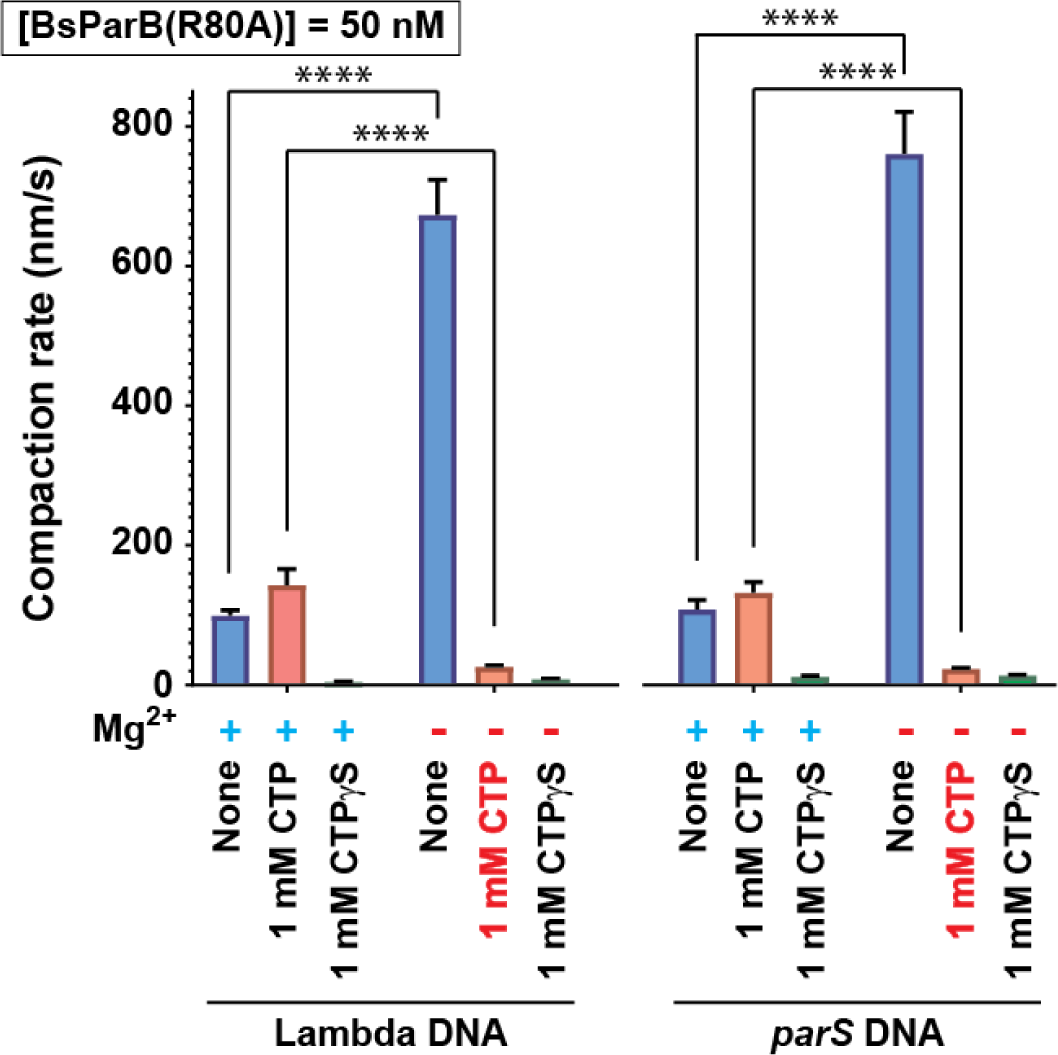
Related to Figures 1 and S3. BsParB(R80A) DNA compaction rates in different conditions. DNA compaction rates by 50 nM BsParB(R80A) on lambda DNA and *parS* DNA in the presence and absence of magnesium ions ([MgCl_2_] = 2.5 mM), CTP (1 mM), and CTPγS (1 mM). DNA Compaction rates in the presence of CTP and in the absence of magnesium ions are highlighted in red. The absence of magnesium and presence of CTP could explain why the previous study23 did not detect DNA compaction by BsParB(R80A). Error bars: s.e.m., **** denotes *p*<0.0001. See Tabs 1 and 2 in the Supplemental File for detailed sample number (*N*) information.

